# Concurrent neuronal activity gates persistent dendritic spine shrinkage during mGluR-induced long-term depression

**DOI:** 10.64898/2026.11.21.765465

**Authors:** E Rojo-Francàs, N Sánchez-Rodríguez, A Santuy, K Hörpudóttir, J Flores, A Parcerisas, E Soriano, A Ranson, K Zito, M Bosch

## Abstract

Synaptic plasticity is a fundamental mechanism underlying learning and memory. Most excitatory synapses reside on dendritic spines, which undergo both functional and structural changes. These two forms of plasticity are typically tightly coupled: spines enlarge during synaptic strengthening (long-term potentiation, LTP) and shrink during synaptic weakening (long-term depression, LTD). While this coupling is well established for NMDA receptor (NMDAR)-dependent LTD, there are conflicting reports concerning whether spine shrinkage is coupled with metabotropic glutamate receptor (mGluR)-dependent LTD.

To investigate the source of these conflicting results, we induced mGluR-LTD with the mGluR1/5 agonist DHPG in organotypic rat hippocampal slice cultures and monitored changes in spine morphology and calcium transients in CA1 pyramidal neurons expressing DsRed2 and GCaMP6.

We found that mGluR activation is accompanied by long-term spine shrinkage only under conditions of elevated hippocampal neuronal activity. This form of plasticity, mGluR-induced neuronal activity-dependent structural LTD (MIND-sLTD), requires mGluR1/5 activation and *de novo* protein synthesis, but not NMDAR signaling. Persistent structural changes occur only when neuronal activity coincides temporally with mGluR activation. MIND-sLTD also occurs at individual synapses: mGluR activation induces persistent single-spine shrinkage only when synchronized with two-photon glutamate uncaging, independently of NMDARs.

These findings identify a novel form of structural plasticity and suggest a coincidence-detection mechanism that integrates mGluR signaling with synaptic activity to consolidate long-term spine remodeling.

## INTRODUCTION

Throughout the nervous system, from neural circuits to individual synapses, function is intimately linked to structure. In the mammalian brain, most excitatory synapses reside on dendritic spines, which provide biochemical compartments for the independent integration of presynaptic inputs (Yuste & Bonhoeffer, 2001). Spine structure is closely correlated with synaptic function: the size of the spine head correlates with the size of the postsynaptic density (PSD), the size of the presynaptic active zone, the number of postsynaptic AMPA-type glutamate receptors (AMPARs), and synaptic strength (Harris & Stevens, 1989; Matsuzaki et al., 2001; Schikorski & Stevens, 1997).

These correlations are not limited to baseline parameters but also extend to their activity-dependent changes. Depending on the pattern of neuronal activity, synapses can undergo lasting increases or decreases in transmission efficiency, known as long-term potentiation (LTP) and long-term depression (LTD), respectively (Malenka & Bear, 2004). At the same time, activity can persistently modify spine morphology (Bosch & Hayashi, 2012). These structural changes often accompany functional changes: LTP induction is associated with spine enlargement (structural LTP, sLTP) (Matsuzaki et al., 2004), whereas LTD induction is commonly associated with spine shrinkage (structural LTD, sLTD) (Nägerl et al., 2004; Oh et al., 2013; K.-I. Okamoto et al., 2004; Q. Zhou et al., 2004). However, under certain conditions, structural and functional plasticity can be dissociated (Sdrulla & Linden, 2007; Thomazeau et al., 2021; X. Wang et al., 2007; Wiegert & Oertner, 2013). This dissociation suggests that structural plasticity may serve biological roles beyond simply mirroring functional changes in synaptic strength.

Together, functional and structural plasticity reshape neural networks and are thought to provide fundamental mechanisms for circuit formation and refinement during development (Stein & Zito, 2019) and information storage during learning and memory (Caroni et al., 2012). Understanding how these two forms of plasticity are coordinated is therefore essential. Although the mechanisms underlying functional plasticity have been extensively studied, less is known about those governing structural plasticity or how the two are linked.

In the hippocampus, the two main forms of LTD can be induced by activation of NMDA-type glutamate receptors (NMDARs) or by activation of group I metabotropic glutamate receptors (mGluRs). Both forms of LTD are expressed through AMPAR internalization (Beattie et al., 2000; Snyder et al., 2001), yet they involve distinct mechanisms and do not occlude one another (Huber et al., 2001). NMDAR-dependent LTD (NMDAR-LTD), induced by low-frequency synaptic stimulation (LFS) (Dudek & Bear, 1992; Oh et al., 2013) or by brief application of the agonist NMDA (Lee et al., 1998), has been widely associated with persistent dendritic spine shrinkage and spine elimination (Bosch & Hayashi, 2012; He et al., 2011; Oh et al., 2013; Q. Zhou et al., 2004). This structural plasticity is triggered by non-ionotropic (metabotropic) NMDAR signaling without calcium influx through the receptor (Stein et al., 2015; Thomazeau et al., 2021).

By comparison, mGluR-dependent LTD (mGluR-LTD) can be induced by paired-pulse low-frequency stimulation (PP-LFS) (Huber et al., 2000) or by brief application of the group I mGluR agonist DHPG (Palmer et al., 1997). Its structural correlates, however, remain unclear, as some studies have reported sustained spine shrinkage following mGluR-LTD induction, whereas others have not. Ramiro-Cortés and Israely (Ramiro-Cortés & Israely, 2013) and Sanderson and colleagues (Sanderson et al., 2022) reported long-lasting spine shrinkage associated with DHPG-induced LTD in organotypic hippocampal slices. In contrast, Vanderklish and Edelman observed spine elongation rather than shrinkage (Vanderklish & Edelman, 2002), and Thomazeau and colleagues detected no structural changes following DHPG-induced LTD in acute hippocampal or cortical slices, or in organotypic hippocampal slices, from rats or mice (Thomazeau et al., 2021). Studies examining spine elimination have likewise reported discrepant observations: while some have detected rapid spine loss (Speranza et al., 2022), others found no effect of a single application of mGluR agonists (Hasegawa et al., 2015; Shinoda et al., 2005). Outside the hippocampus, Sdrulla and Linden similarly found no spine structural changes in Purkinje cells after induction of mGluR1-dependent cerebellar LTD (Sdrulla & Linden, 2007).

These contrasting findings raise the question of what determines whether mGluR-LTD is accompanied by spine shrinkage. Here, we sought to identify the conditions and mechanisms that govern this structural response. We found that long-term spine shrinkage associated with DHPG-induced LTD occurs only when concomitant neuronal activity in the hippocampal network is present. This form of plasticity requires mGluR1/5 activation and *de novo* protein synthesis, but not necessarily NMDARs. Using glutamate uncaging, we further observed a local activity requirement at individual synapses. These findings identify a form of structural plasticity in which mGluR signaling converge with synaptic activity to consolidate spine structural remodeling.

## RESULTS

### Persistent structural plasticity of dendritic spines associated with mGluR-induced LTD depends on concurrent neuronal activity

To carry out a longitudinal study of the structural changes in dendritic spines associated with the induction of mGluR-LTD, we used organotypic hippocampal slice cultures prepared from rats at postnatal days 6-8 (P6-8). We biolistically transfected the slices with a plasmid expressing DsRed2 as a volume marker to observe spine and neuron morphology in well-isolated neurons, together with a plasmid expressing the genetically encoded calcium sensor GCaMP6s to simultaneously monitor the ongoing neuronal activity in the hippocampal network (**Fig. 1A-C**). We performed time-lapse live imaging using a fluorescence confocal microscope while continuously perfusing the slices with carbogenated artificial cerebrospinal fluid (ACSF) at 32 °C. We selected CA1 pyramidal neurons and monitored a subpopulation of their spines on the apical dendrites in the *stratum radiatum* (**Fig. 1D-E**) for over 92 min. We induced chemical mGluR-LTD by brief (10 min) bath application of the group I mGluR agonist (R,S)-3,5-dihydroxyphenylglycine (DHPG, 100 μM).

**Figure 1.**
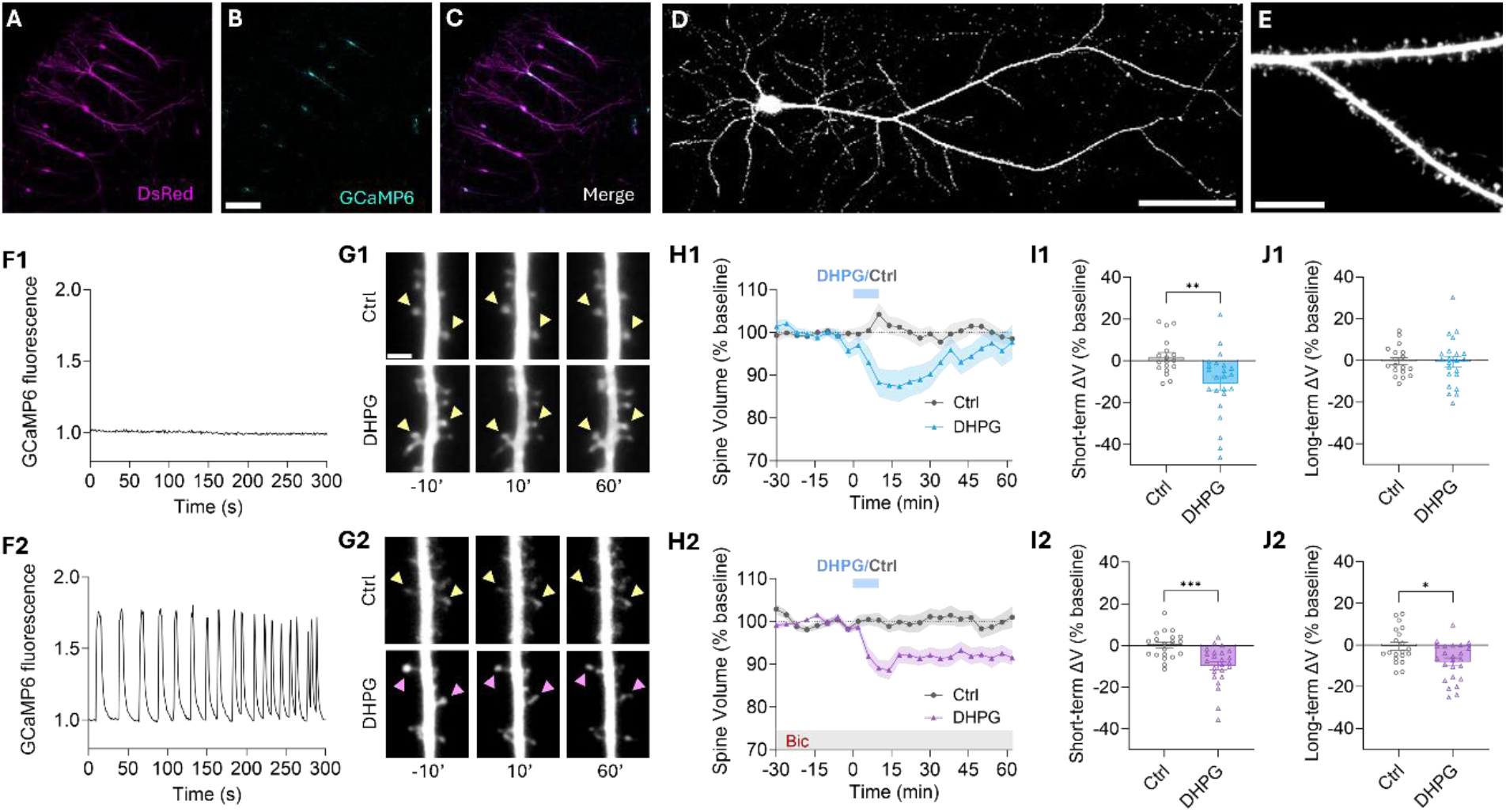
Induction of LTD through activation of mGluRs with DHPG is associated with persistent dendritic spine shrinkage only when neuronal network activity is present. **A-C)** Confocal images of the CA1 region of an organotypic rat hippocampal slice showing neurons expressing DsRed2 (A), GCaMP6s (B) and the merged image of both channels (C). Scale bar represents 200 µm. **D-E)** Confocal images of a CA1 pyramidal neuron expressing DsRed2 (D) and a magnified view of dendrites with dendritic spines (E). Scale bars represent 100 µm and 10 µm, respectively. **F)** Time course of neuronal activity levels in the hippocampal slice detected by calcium imaging (GaCMP6 fluorescence signal) in the absence (F1) or presence of 10 μM bicuculline (Bic) (F2). **G)** Representative images of dendritic spines 10 min before and 10 min and 60 min after mock (Ctrl) or DHPG application, in the absence (G1) or presence of Bic (G2). Scale bar, 2 µm. **H)** Time course (in min) of mean spine volume ± SEM (% of baseline) during bath application of 100 µM DHPG for 10 min (blue bar) or no application (control), in the absence (H1) or presence of Bic (H2). **I-J)** Quantification of spine volume changes (% relative to baseline, with negative values indicating a decrease and positive values an increase) during the 6-18 min period after DHPG application onset (I, short-term structural depression, sSTD) and during the 46-62 min period after DHPG (J, long-term structural depression, sLTD) under control (gray) and DHPG (blue or violet) conditions. Values are presented as mean ± SEM; each point represents one dendrite. In the absence of Bic, significant short-term spine shrinkage is observed compared with the control condition (I1; control: 1.86 ± 2.08 %, n = 18; DHPG: -10.91 ± 3.29 %, n = 23), but it is not maintained in the long term (J1; control: -0.42 ± 1.70 %, n = 18; DHPG: -0.84 ± 2.48 %, n = 23). In the presence of Bic, significant short-term spine shrinkage is observed compared with the control condition (I2; control: +0.14 ± 1.45 %, n = 20; DHPG: -9.70 ± 1.91 %, n = 24) and is maintained in the long term (J2; control: -0.56 ± 1.88 %, n = 20; DHPG: -8.04 ± 1.84 %, n = 24). Statistical significance was assessed using an unpaired t-test with Welch’s correction for I1 and J1, a Mann–Whitney test for I2, and an unpaired t-test for J2. ***p < 0.001, **p < 0.01, *p < 0.05. n = number of dendrites.

In a previous study, we observed that induction of mGluR-LTD in hippocampal slices was not accompanied by spine shrinkage, even though DHPG triggered a reduction in synaptic strength (i.e., functional LTD) and internalization of AMPARs (Thomazeau et al., 2021). However, since other studies observed persistent spine shrinkage during mGluR-LTD using similar experimental approaches (Ramiro-Cortés & Israely, 2013; Sanderson et al., 2022), we postulated that an additional uncontrolled parameter might determine the structural fate of spines during mGluR-LTD. We hypothesized that this parameter could be the endogenous levels of network neuronal activity in the hippocampal slice (Thomazeau et al., 2021), which can vary across experiments and laboratories depending on the experimental conditions. To explore this idea, we analyzed the levels of spontaneous neuronal activity in the hippocampal slices before LTD induction using the GCaMP6 fluorescence signal. We observed that the basal activity levels in our slices were very low (**Fig. 1F1**). In these slices, application of DHPG resulted in transient short-term spine shrinkage (structural short-term depression, sSTD; an average spine volume reduction of 10.91 ± 3.29 % relative to baseline, i.e. ΔV = - 10.91 ± 3.29 %, during the 6-18 min period after DHPG application; **Fig. 1G1-H1**), which was significantly different from that in control experiments (control, no DHPG application, **Fig. 1I1**). However, this shrinkage was not persistent, and spines returned to their original average size (a spine volume reduction of 0.84 ± 2.48 % relative to baseline, i.e. ΔV = - 0.84 ± 2.48 %, during the 46-62 min period after DHPG application, **Fig. 1G1-H1**). Thus, similar to our previous study (Thomazeau et al., 2021), we did not detect significant long-term spine shrinkage (structural long-term depression, sLTD, **Fig. 1J1**) after mGluR activation.

To increase basal hippocampal activity, we applied the GABAA receptor antagonist bicuculline (Bic). Blocking the inhibitory input in hippocampal slices results in the generation of spontaneous highly synchronized epileptiform activity bursts (Miles & Wong, 1983). Application of 10 μM bicuculline to our slices readily induced a synchronized rhythmic pattern of neuronal activity with an average of 3.90 ± 0.82 calcium peaks/min (**Fig. 1F2**). Bicuculline was applied 1 h before DHPG addition and maintained throughout the experiment. Under these conditions, we detected both short-term (ΔV = - 9.7 ± 1.91 %) and long-term spine shrinkage (ΔV = -8.04 ± 1.84 %) after mGluR-LTD induction. This shrinkage persisted for at least 1 hour (**Fig. 1G2-H2**) and was significantly different from that under control conditions (**Fig. 1J2**). We conclude that mGluR-LTD induces persistent spine shrinkage only when highly synchronized neuronal activity is present in the hippocampal circuit.

**Figure 2.**
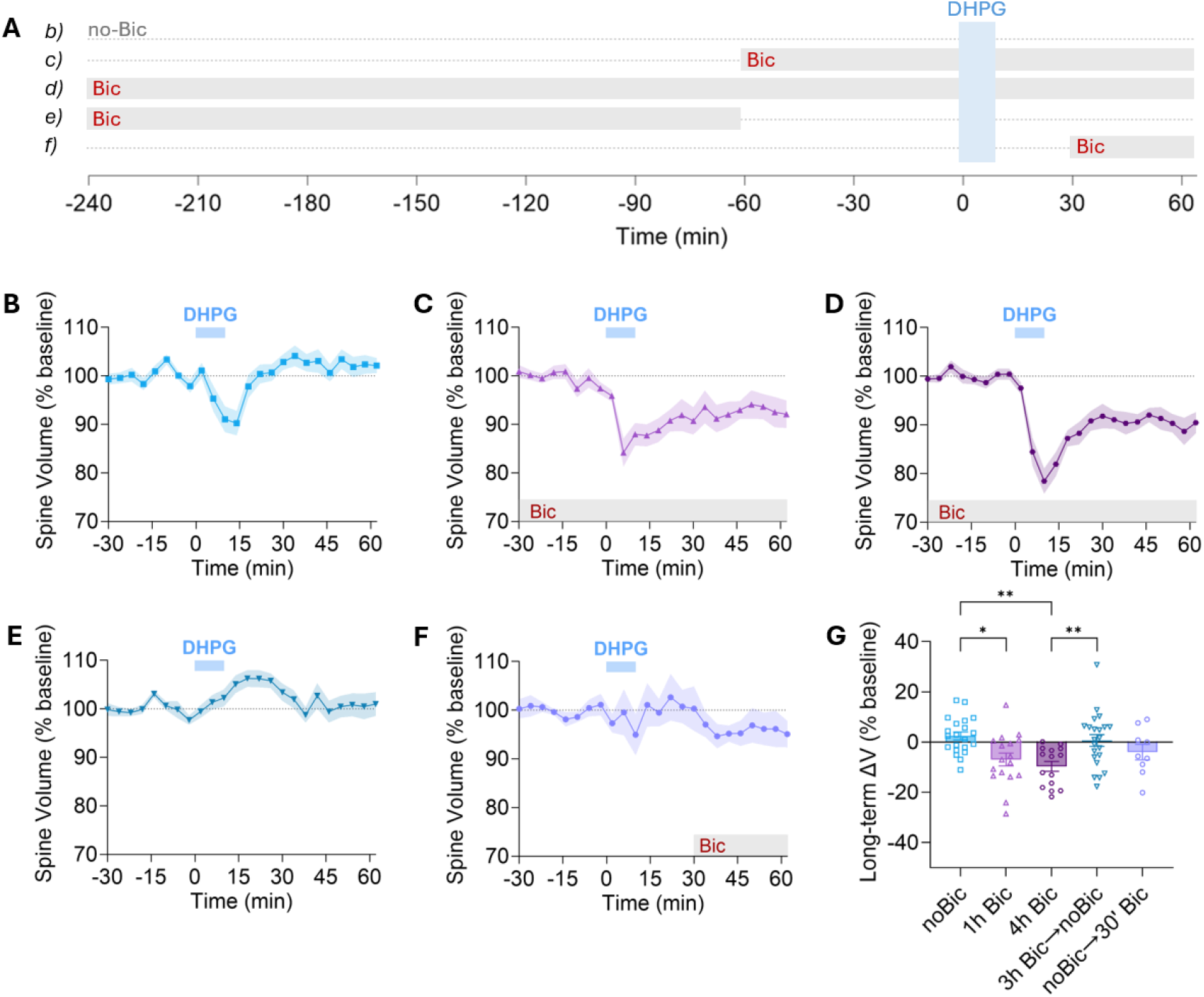
Persistent spine shrinkage occurs only when mGluR activation and neuronal activity are temporally coincident. **A)** Schematic representation of the experimental conditions, in which slices were exposed to 10 μM bicuculline (Bic, gray bars) for different periods relative to the application of 100 μM DHPG (10 min, blue bar): no Bic exposure (b), Bic exposure starting 1 h (c), or 4 h (d) before DHPG application and maintained throughout the experiment, Bic exposure for 3 h followed by withdrawal 1 h before DHPG (e), and Bic exposure starting 30 min after DHPG application onset (f). **B-F)** Time course (in min) of mean spine volume (% of baseline) ± SEM under the different Bic treatments. DHPG application in the absence of Bic (B) did not induce persistent spine shrinkage (noBic, ΔV = +2.48 ± 1.57 %, n = 21 dendrites). Bic treatment for 1 h (C) resulted in DHPG-induced persistent spine shrinkage (1h Bic, ΔV = -6.92 ± 2.55 %, n = 17), as did Bic treatment for 4 h (D, 4 h Bic, ΔV = -9.66 ± 2.01 %, n = 15). In contrast, Bic preincubation for 3 h followed by withdrawal 1 h before DHPG application (E) did not produce spine shrinkage (3h Bic->noBic, ΔV = +0.69 ± 2.33 %, n = 22). Similarly, delayed Bic exposure 30 min after DHPG application (F) failed to induce spine shrinkage (noBic->30’ Bic, ΔV = -3.94 ± 3.12 %, n = 9). **G)** Quantification of spine volume change (% relative to baseline) during the last 16 min of imaging (long-term shrinkage, sLTD). Persistent spine shrinkage was observed only under conditions in which bicuculline was present during the DHPG application. Values are presented as mean ± SEM; each point represents one dendrite. Statistical significance was determined using one-way ANOVA with post-hoc Tuckey correction. ** p < 0.01, * p < 0.05.

### mGluR activation and neuronal activity must be temporally coincident to induce persistent spine shrinkage

We next sought to characterize the temporal requirements for this form of structural plasticity, which we term mGluR-induced neuronal activity-dependent structural LTD (MIND-sLTD). We wanted to determine whether neuronal activity had to strictly coincide temporally with mGluR activation, whether prior neuronal activity could prime spines to shrink in response to mGluR activation, or, conversely, whether DHPG could prime spines to shrink in response to subsequent neuronal activity. To address these possibilities, we compared 5 conditions in which bicuculline was added for different durations, either before, during or after DHPG application (**Fig. 2A**). In every experiment in which bicuculline was added, we performed calcium imaging to confirm the presence of the characteristic pattern of synchronized neuronal activity in the entire hippocampal slice and, specifically, in the neuron whose spines were examined.

We again confirmed that, in the absence of bicuculline, there was no persistent spine shrinkage after DHPG application (ΔV = +2.48 ± 1.57 %; **Fig. 2B**), whereas 1 h of pre-exposure to bicuculline resulted in sustained spine shrinkage (ΔV = -6.92 ± 2.55 %; **Fig. 2C**). We next tested whether a longer exposure to neuronal activity resulted in a greater degree of MIND-sLTD, i.e., more pronounced spine shrinkage. Longer exposure to neuronal activity might increase activity-induced synthesis of plasticity-related proteins, such as Arc/Arg3.1, which is strongly implicated in the expression of functional mGluR-LTD (Newpher et al., 2018; Park et al., 2008; Waung et al., 2008). We extended bicuculline exposure to 4 h before DHPG application (**Fig. 2D**) to allow more time for new protein synthesis. Under this condition, we observed a trend toward more pronounced short-term (ΔV = -16.76 ± 1.99 %) and long-term spine shrinkage (ΔV = - 9.66 ± 2.01 %; **Fig. 2D, G**) but neither significantly differed from that observed after 1 h of bicuculline exposure.

Next, to assess whether neuronal activity could prime spines for mGluR-induced shrinkage, we exposed the slices to bicuculline for 3 h and then withdrew it 1 hour before DHPG application (**Fig. 2E**). We confirmed that the synchronized activity pattern had completely disappeared by 30 min before DHPG application. Under this condition, MIND-sLTD did not occur, as we did not observe any short-term (ΔV = +3.73 ± 1.42 %) or long-term spine shrinkage (ΔV = +0.69 ± 2.33 %; **Fig. 2G**). These results argue against the possibility that prior activity-induced synthesis of plasticity-related proteins is sufficient to induce spine shrinkage following mGluR activation. Finally, to assess the opposite possibility, that mGluR activation primes spines to shrink in response to subsequent neuronal activity, we exposed slices to bicuculline 30 min after DHPG (**Fig. 2F**). Under this condition, MIND-sLTD did not occur either, as we did not observe any short-term (ΔV = -1.50 ± 4.33 %) or any significant long-term spine shrinkage (ΔV = -3.94 ± 3.12 %). Altogether, these experiments suggest that mGluR activation and neuronal activity do not prime each other but must be temporally coincident to produce persistent shrinkage of dendritic spines.

### Molecular mechanisms of mGluR-induced neuronal activity-dependent structural LTD (MIND-sLTD)

We next aimed to dissect the molecular and signaling mechanisms underlying this form of structural plasticity. We repeated the same experiment, activating mGluRs with DHPG for 10 min during constant exposure to bicuculline (applied 1 hour before DHPG and maintained in the bath throughout the experiment, as in Fig. 1H2 and Fig. 2C), and tested the effects of inhibiting selected signaling elements involved in synaptic plasticity (**Fig. 3A**). We first confirmed that spine shrinkage was directly induced by the activation of group I mGluRs by applying the mGluR1-selective antagonist LY367385 and the mGluR5-selective antagonist MTEP (Volk et al., 2006) (**Fig. 3B**). Under these conditions, both the initial and sustained phases of spine shrinkage were completely abolished, i.e., no detectable changes in spine volume were observed at any time point (**Fig. 3F-G**). Next, to confirm the role of neuronal activity in gating mGluR-induced spine shrinkage, we blocked action potentials in the entire hippocampal slice using the voltage-gated sodium channel blocker tetrodotoxin (TTX; **Fig. 3C**). We confirmed that calcium oscillations were completely absent in the slice and in the neuron under examination. Under these conditions, we did not observe any spine shrinkage following DHPG application, either short-term or long-term (**Fig. 3F-G**), which is consistent with the study by Ramiro-Cortés and Israely (Ramiro-Cortés & Israely, 2013). These results confirm that MIND-sLTD requires both group I mGluR activation and the presence of network neuronal activity to be expressed.

**Figure 3.**
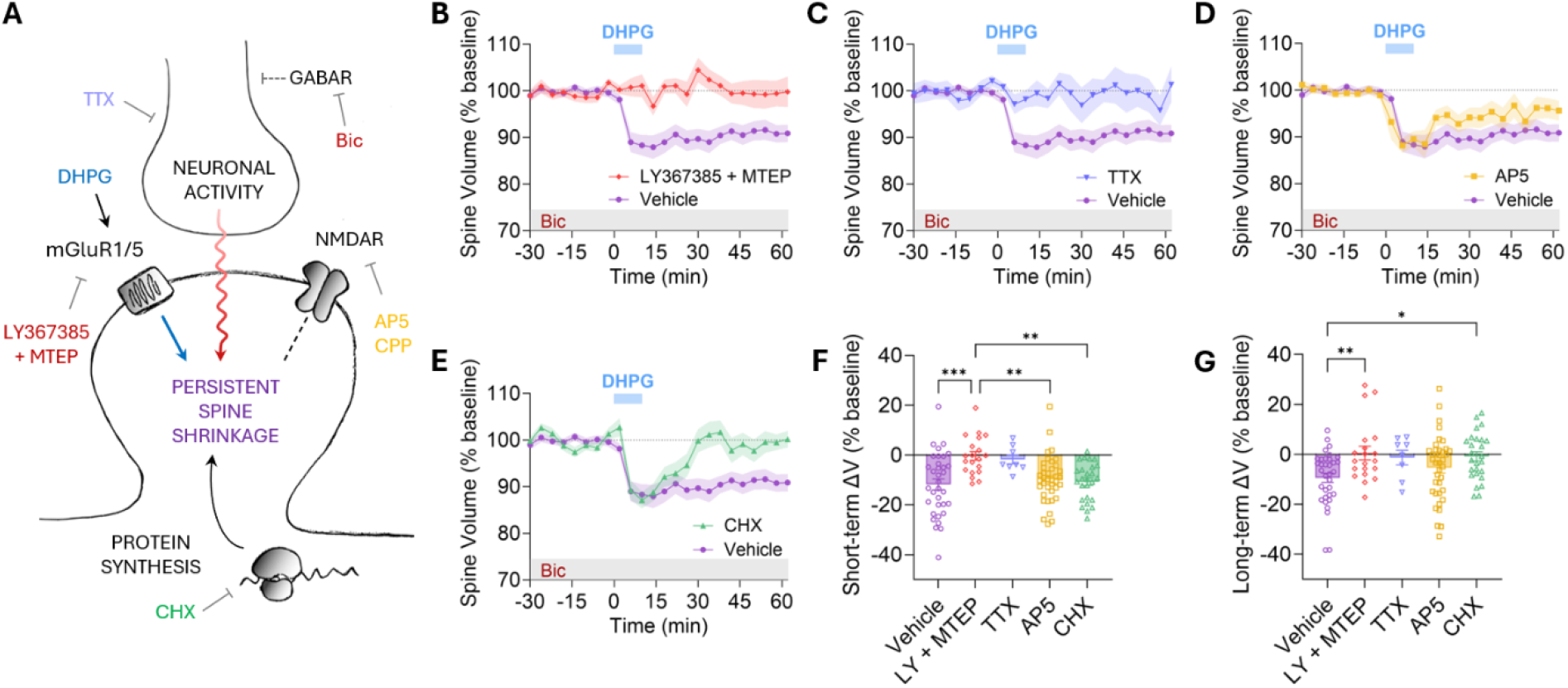
mGluR-induced neuronal activity-dependent structural LTD (MIND-sLTD) depends on mGluR1/5, action potentials, and new protein synthesis, but not on NMDARs. **A)** Schematic model of the mechanism underlying MIND-sLTD in hippocampal dendritic spines. **B-E)** Time course (in min) of mean spine volume (% of baseline) ± SEM during 100 μM DHPG application (10 min, blue bar) in the presence of 10 μM bicuculline (Bic, gray bar) under control conditions (vehicle) or in the presence of the following pharmacological agents: the mGluR1 antagonist LY367385 (50 µM) and the mGluR5 antagonist MTEP (1 µM, B), the voltage-gated sodium channel blocker tetrodotoxin (TTX, 1 µM) (C), the competitive NMDAR antagonist AP5 (50 µM, D), or the protein synthesis inhibitor cycloheximide (CHX, 60 µM) (E). **F-G)** Quantification of spine volume changes (% relative to baseline) during the 6-18 min period after DHPG application onset (F, short-term spine shrinkage, sSTD) and during the 46-62 min period after DHPG (G, long-term spine shrinkage, sLTD) under control (vehicle) and pharmacological treatments. LY367385 and MTEP prevented both DHPG-induced sSTD (vehicle: ΔV = -11.80 ± 2.16 %; LY + MTEP: ΔV = -0.34 ± 1.73 %) and sLTD (vehicle: ΔV = -9.35 ± 1.87 %, n = 34 dendrites; LY + MTEP: ΔV = 0.55 ± 2.72 %, n = 20 dendrites). TTX showed a tendency to block both DHPG-induced sSTD (ΔV = -1.72 ± 1.84 %) and sLTD (ΔV = -1.26 ± 2.93 %, n = 8). AP5 did not prevent DHPG-induced sSTD (ΔV = - 9.69 ± 1.52 %) and did not significantly prevent sLTD (ΔV = -5.29 ± 2.11 %, n = 38). CHX did not prevent DHPG-induced sSTD (ΔV = -10.65 ± 1.49 %) but completely blocked sLTD (ΔV = -0.83 ± 1.84 %, n = 26). Values are presented as mean ± SEM; each point represents one dendrite. Statistical significance was determined using one-way ANOVA with post-hoc Tuckey correction. *** p < 0.001 ** p < 0.01, * p < 0.05.

Multiple studies have established crosstalk between mGluRs and NMDARs in regulating synaptic transmission, synaptic plasticity (Collingridge et al., 2004) and structural plasticity (Oh et al., 2013). Activation of mGluRs affects NMDAR responses (Attucci et al., 2001; Mannaioni et al., 2001; Skeberdis et al., 2001) and activation of NMDARs affects mGluR responses (Alagarsamy et al., 1999; Palmer et al., 1997; Sanderson et al., 2022). Hence, we next investigated the specific role that NMDARs play in MIND-sLTD. We used the competitive antagonist AP5 to directly block NMDARs during mGluR activation with DHPG in the presence of bicuculline (**Fig. 3D**). We found that AP5 had no effect on the transient phase of spine shrinkage (**Fig. 3F**) but we detected a non-significant trend toward a partial suppression of the persistent phase (ΔV = - 5.29 ± 2.11 % with AP5 versus ΔV = *- 9.35 ± 1.87 %* with vehicle, p = 0.2929; **Fig. 3G**). However, we still observed a significant reduction in spine volume (ΔV = - 5.29 ± 2.11 %, p = 0.033 compared with baseline, paired t-test), indicating that persistent spine shrinkage still occurs during NMDAR blockade. The high degree of variability between experiments in AP5 treatment, especially during the persistent phase (**Fig. 3G**), may indicate the presence of yet another uncontrolled parameter that determines whether NMDARs contribute to the mGluR-induced spine shrinkage. In any case, our results suggest that NMDARs are not necessary for the expression of MIND-sLTD, which is consistent with the reports of Ramiro-Cortés and colleagues (Ramiro-Cortés et al., 2026; Ramiro-Cortés & Israely, 2013).

We finally assessed the involvement of protein synthesis in the expression of MIND-sLTD. Many forms of synaptic plasticity depend on the synthesis of new proteins, including forms of functional plasticity such as mGluR-LTD (Huber et al., 2000) and the late phases of NMDAR-LTP and NMDAR-LTD (Pfeiffer & Huber, 2006). Some forms of structural plasticity also require new protein synthesis, including NMDAR-sLTD (Stein et al., 2015; Thomazeau et al., 2021), the late phase of NMDAR-sLTP (Bosch et al., 2014; Tanaka et al., 2008) and, interestingly, both early and late phases of mGluR-sLTD, as seen in those studies detecting spine shrinkage (Ramiro-Cortés et al., 2026; Ramiro-Cortés & Israely, 2013). We, therefore, asked whether MIND-sLTD was similarly dependent on *de novo* protein synthesis. We applied the translation inhibitor cycloheximide (CHX; **Fig. 3E**) and found that, while the short-term spine shrinkage following DHPG application was not altered (**Fig. 3F**), the long-term spine shrinkage was abolished (**Fig. 3G**) and spines recovered their original size ∼30 min after DHPG application. Thus, MIND-sLTD does not require synthesis of new proteins to be initially induced but requires *de novo* translation for persistent consolidation.

Altogether, our results point to a molecular mechanism in which MIND-sLTD requires mGluR1/5 activation, the presence of action potentials in the circuit, and the synthesis of new proteins for its consolidation, but does not strictly require NMDAR activation.

### MIND-sLTD occurs at the level of single spines

We finally sought to study this phenomenon at the maximum temporal and spatial resolution: the individual synapse. The experiments described thus far relied on global pharmacological manipulations that affect the entire neuronal network. These approaches demonstrate that high levels of neuronal activity are required for mGluR-induced persistent spine shrinkage, but do not resolve whether activity must occur locally at individual synapses or globally across the network.

We induced MIND-sLTD at a single synapse by combining a pharmacological approach to globally activate mGluRs, using bath application of DHPG, with focal photostimulation to locally mimic synaptic activity, using two-photon photolysis of caged glutamate at the target spine (**Fig. 4A**). Single-spine NMDAR-sLTD has been achieved in the past using two-photon low-frequency glutamate uncaging (LFU) (Oh et al., 2013; Jang et al., 2021), and single-spine mGluR-LTD has also been achieved using two-photon LFU in the presence of NMDAR inhibitors (Holbro et al., 2009). Holbro and colleagues, however, did not report any associated structural changes in spines after mGluR-LTD. A very recent study developed a paired-pulse LFU (ppLFU) protocol to induce single-spine mGluR-LTD and reported long-lasting spine shrinkage (Ramiro-Cortés et al., 2026). Here, we combined glutamate uncaging to mimic neuronal activity at a single synapse with pharmacological activation of group I mGluRs. Our uncaging protocol (0.4 ms pulses of 720 nm at 0.1 Hz for 10 min), adapted from previous work (Jang et al., 2021) and termed subthreshold low-frequency uncaging (subLFU; **Fig. 4B**), did not induce any change in spine volume by itself, either growth or shrinkage.

**Figure 4.**
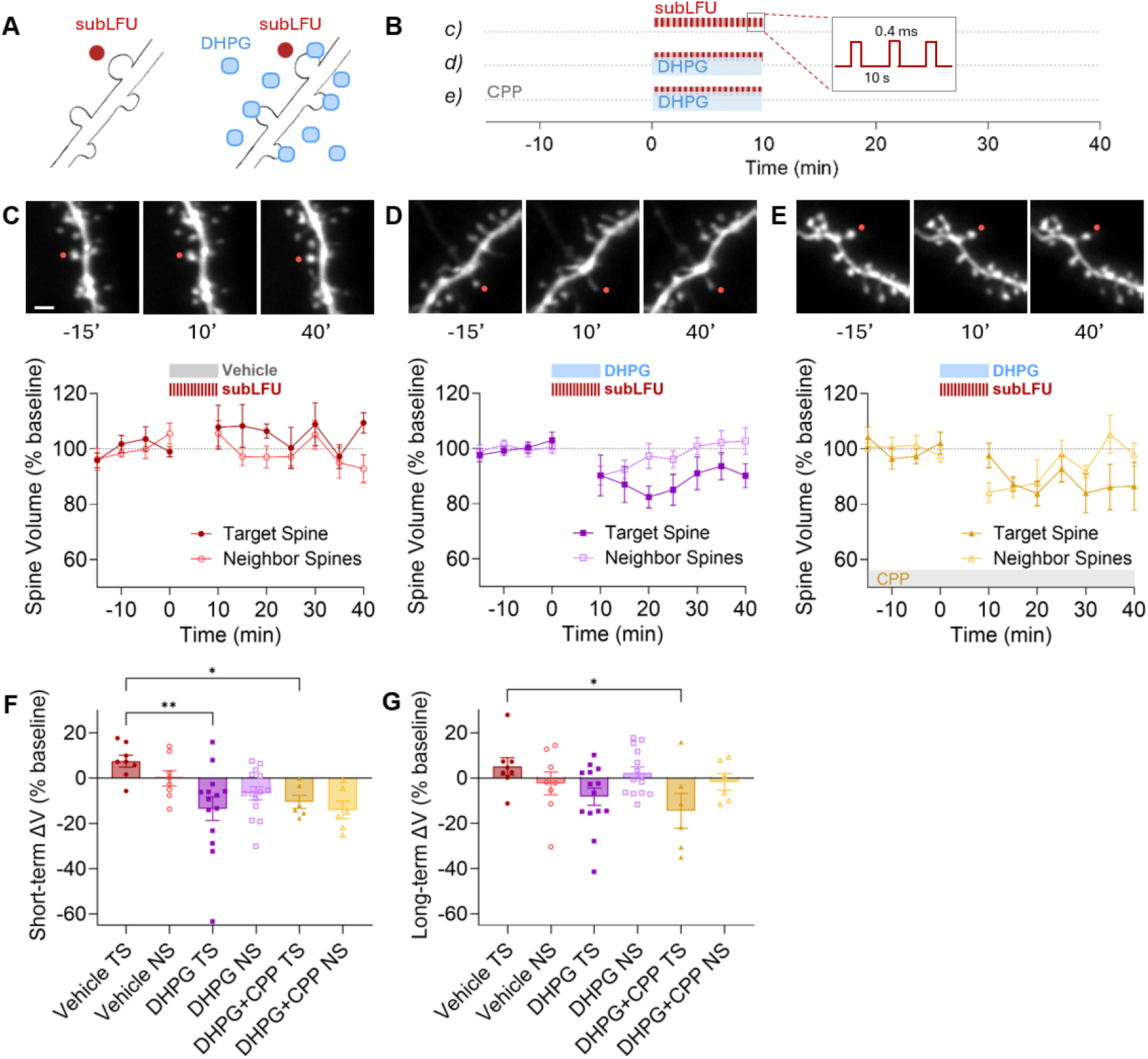
Concurrent mGluR activation and glutamate uncaging-induced synaptic activity gate persistent shrinkage of single spines. **A)** Schematic of the experimental approach. An individual target dendritic spine was stimulated (red dot) using a subthreshold low-frequency uncaging (subLFU) protocol, with or without DHPG bath application (blue dots). **B)** Schematic representation of the experimental protocol and conditions. Target spines were stimulated using the subLFU protocol (0.4-ms pulses of 720 nm pulsed laser delivered at 0.1 Hz for 10 min), while the entire slice was treated with vehicle (c) or DHPG (d), in the absence (c, d) or presence (e) of the competitive NMDAR antagonist CPP. **C-E)** Representative images of a glutamate uncaging-stimulated target spine (red dot) and non-stimulated neighboring spines (scale bar, 2 *µm), together with the corresponding time courses (in min) of mean spine volume ± SEM (% of baseline) following subLFU stimulation (10 min) in combination with vehicle application (C), 100 μM DHPG bath application (D) or 100 μM DHPG bath application in the presence of CPP (E). **F)** Quantification of spine volume changes (% relative to baseline) during the 10-15 min interval after subLFU stimulus onset (short-term shrinkage, sSTD) in stimulated target spines (TS) and neighboring unstimulated spines (NS). Target spines receiving subLFU stimulation together with DHPG exhibited a significant reduction in spine volume compared with vehicle-treated controls, both in the presence and absence of CPP, indicating that the initial structural response does not require NMDAR signaling (Vehicle TS: ΔV = +7.48 ± 2.65 %, n = 8; DHPG TS: ΔV = - 13.47 ± 5.15 %, n = 14; DHPG + CPP TS: ΔV = -10.44 ± 2.88 %, n = 6). **G)** Quantification of spine volume changes (% relative to baseline) during the 30-40 min interval after subLFU stimulus onset (long-term shrinkage, sLTD). Stimulated target spines showed a trend toward a persistent reduction in spine volume following combined subLFU and DHPG stimulation. Significant persistent shrinkage was observed in stimulated spines in the presence of CPP, supporting an NMDAR-independent mechanism of MIND-sLTD at single synapses (Vehicle TS: ΔV = +5.13 ± 3.86 %, n = 8; DHPG TS: ΔV = -8.24 ± 3.81 %, n = 14; DHPG + CPP TS: ΔV = -14.45 ± 7.70 %, n = 6). Values are presented as mean ± SEM; each point represents one spine. Statistical significance for short-term ΔV (sSTD) was determined using a Kruskal-Wallis test with post-hoc Dunn’s correction. Statistical significance for long-term ΔV (sLTD) was determined using one-way ANOVA test with post-hoc Bonferroni correction. * p < 0.05, ** p < 0.01. n = number of spines*.

We used this neutral stimulus protocol to test the effect of combining local synaptic activity with mGluR activation on the structural plasticity of an individual spine. These experiments were performed in mouse hippocampal slice cultures, biolistically transfected with EGFP. TTX was included in the ACSF to block circuit action potentials, permitting us to test whether local glutamatergic activity was sufficient to induce MIND-sLTD. We selected a target spine and delivered the subLFU stimulation protocol for 10 min (**Fig. 4B**). Under control conditions, in which we administered vehicle instead of DHPG (**Fig. 4C**), the subLFU stimulation alone did not produce any structural change either in the short term (no sSTP or sSTD in the 10-15 min interval after vehicle application compared to baseline) or in the long term (no sLTP or sLTD during the 30-40 min interval after vehicle application), in either the two-photon laser-targeted spines or in the neighboring spines. However, when 100 µM DHPG was applied simultaneously with subLFU stimulation (**Fig. 4D**), the targeted spine showed significant short-term spine shrinkage (sSTD, ΔV = - 13.47 ± 5.15 % of baseline; p = 0.014 compared with baseline, paired t-test; p = 0.004 compared with vehicle-treated spines, Kruskal-Wallis test; **Fig. 4F**). This shrinkage persisted in the long term (sLTD, ΔV = - 8.24 ± 3.81 % of baseline; p = 0.047 compared with baseline, paired t-test; p = 0.032 compared with vehicle-treated spines, one-way ANOVA). However, the comparison with vehicle-treated spines did not reach significance when all groups, including those from the CPP experiments in Fig. 4E, were included in the analysis (p = 0.144, one-way ANOVA; **Fig. 4G**). We also observed a transient decrease in spine volume in all non-targeted neighboring spines treated with DHPG (p = 0.031 compared with their baseline, paired t-test, **Fig. 4D**), but this shrinkage was not persistent as spines quickly recovered their baseline volume (**Fig. 4G**). This short-term effect was consistent with our previous results, in which global DHPG application induced short-term spine shrinkage that was not consolidated in the long term in the absence of neuronal activity (**Fig. 1G1**, **Fig. 2B**).

These results suggest that local synaptic activity can stabilize the mGluR-induced structural plasticity at an individual synapse. They indicate that global network neuronal activity is not strictly required but that the homosynaptic coincidence between mGluR activation and synaptic activity is sufficient. Thus, we conclude that the phenomenon of MIND-sLTD also occurs at the level of a single synapse. These results also narrow the temporal requirements for MIND-sLTD: activity does not need to be present long before or after mGluR activation but can be restricted to the same 10-min period, which refines the temporal coincidence requirement identified in the pharmacological experiments in Fig. 2.

Finally, we assessed the role that NMDARs might play in the expression of single-spine MIND-sLTD. In our previous experiments, we showed that inhibition of NMDARs with AP5 did not block the transient spine shrinkage and persistent shrinkage remained detectable after MIND-sLTD induction (**Fig. 3D**). Here, we used the competitive antagonist CPP to similarly block NMDARs. We found that CPP did not abolish the DHPG-induced spine shrinkage in subLFU-targeted spines (**Fig. 4E**), either in the short term (p = 0.022 compared with vehicle-treated spines, Kruskal-Wallis test with multiple comparisons across all groups; **Fig. 4F**) or in the long term (p = 0.043 compared with vehicle-treated spines, one-way ANOVA with multiple comparisons across all groups; **Fig. 4G**). We also observed transient spine shrinkage across neighboring spines (p = 0.010 compared with their baseline, paired t-test, **Fig. 4E-F**), which did not persist in the long term (**Fig. 4E, G**), as expected after DHPG application in the absence of neuronal or synaptic activity. These results demonstrate that MIND-sLTD is NMDAR-independent at the level of single spines.

Altogether, our findings demonstrate that induction of mGluR-LTD is not accompanied by persistent shrinkage of dendritic spines unless temporally coincident synaptic activity is present. Our results suggest that this novel form of structural plasticity requires *de novo* protein synthesis but not necessarily NMDAR activation for its consolidation.

## DISCUSSION

The main goal of this study was to investigate the structural plasticity of dendritic spines associated with mGluR-LTD induction and find a possible explanation for the discrepant structural outcomes reported in the literature. We wanted to identify the conditions under which mGluR activation translates into long-lasting structural remodeling of dendritic spines. We found that temporally coincident neuronal activity permitted persistent mGluR-induced spine shrinkage. This persistence required new protein synthesis but not NMDAR signaling. This phenomenon, which we term mGluR-induced neuronal activity-dependent structural LTD (MIND-sLTD), was reproduced at the level of individual spines by combining global DHPG application with local glutamate uncaging.

### Concurrent neuronal activity gates persistent spine shrinkage and explains the controversy in mGluR-sLTD

Although mGluR-LTD has been extensively characterized at the electrophysiological level (Gladding et al., 2009), its structural correlates remain less well understood (Bosch & Hayashi, 2012). Studies addressing this question have reported markedly different outcomes. Ramiro-Cortés and Israely investigated DHPG-induced LTD in mouse organotypic hippocampal slices and found long-lasting (up to 3 h) spine shrinkage (Ramiro-Cortés & Israely, 2013). Similarly, Sanderson and colleagues studied DHPG-induced LTD in rat organotypic hippocampal slices and detected sustained (up to 30 min) spine shrinkage (Sanderson et al., 2022). However, other studies using similar approaches did not observe such structural changes. Vanderklish and Edelman observed spine elongation 30 min after DHPG in both rat dissociated hippocampal neuronal cultures and rat organotypic hippocampal slices, although they did not explicitly measure volume changes (Vanderklish & Edelman, 2002). Holbro and colleagues were the first to induce single-spine mGluR-LTD using glutamate uncaging (Holbro et al., 2009) but did not report any associated structural response. Finally, Thomazeau and colleagues specifically addressed the structural correlates of both NMDAR-LTD and mGluR-LTD in a variety of preparations, including acute and cultured hippocampal and cortical slices, from mice and rats. Although they detected robust persistent spine shrinkage after NMDAR-LTD induction, they failed to detect any transient or persistent spine shrinkage after mGluR-LTD induction in any of these preparations (Thomazeau et al., 2021). Similarly, studies examining spine elimination after mGluR-LTD have reported discrepant results, with one study reporting rapid and robust spine elimination 30 min after DHPG application in hippocampal slices and dissociated cultures (Speranza et al., 2022), while others reported that a single application of mGluR agonists had no effect, but repeated applications on consecutive days produced spine loss (Hasegawa et al., 2015; Shinoda et al., 2005). In the cerebellum, Sdrulla and Linden observed a dissociation between functional and structural plasticity of dendritic spines in Purkinje cells: induction of mGluR1-dependent LTD, although mechanistically different from hippocampal mGluR-LTD, was not accompanied by spine shrinkage (Sdrulla & Linden, 2007).

Taking all these discrepant reports into consideration, we hypothesized that some experimental variable might be responsible for these variations in structural outcome. To investigate this question, we chose organotypic hippocampal slice cultures from rats and mice as they recapitulate the major structural and functional features of the intact hippocampus, including dendrite and spine maturation (De Simoni et al., 2003), and spontaneous and evoked patterns of network activity (K. Okamoto et al., 2014), while allowing easy genetic and pharmacological manipulation. Importantly, this approach facilitated comparison with previous studies reporting different structural outcomes using similar experimental approaches.

We had previously hypothesized that neuronal activity could explain the controversial results in mGluR-sLTD reported in the literature (Thomazeau et al., 2021). To test this hypothesis, we used the GABAAR inhibitor bicuculline to block the inhibitory inputs in the hippocampal circuitry, which results in the generation of spontaneous highly synchronous epileptiform activity oscillations (Miles & Wong, 1983; K. Okamoto et al., 2014). Under these conditions, we detected a sustained phase of spine shrinkage that persisted for at least 1 h (Figs. 1-3). Few studies have addressed the effect that disinhibition has on the expression of mGluR-LTD and, again, they yielded conflicting results: while Palmer and colleagues found that picrotoxin enhanced mGluR-LTD in an NMDAR-independent way (Palmer et al., 1997), Rohde and colleagues observed no alteration of the magnitude of functional mGluR-LTD in the presence of gabazine (Rohde et al., 2009).

Nonetheless, here we studied the effects of disinhibition on structural, not functional LTD, which has been unexplored to our knowledge. In this case, disinhibition with bicuculline did make a difference, as it converted a transient mGluR-sSTD into a persistent mGluR-sLTD. We therefore propose that differences in intrinsic circuit activity in the hippocampal preparations represent a plausible explanation for at least some of the discrepancies reported in the literature. It is possible that those studies that observed spine shrinkage after induction of hippocampal mGluR-LTD (Ramiro-Cortés et al., 2026; Ramiro-Cortés & Israely, 2013; Sanderson et al., 2022) had high levels of spontaneous activity, while those that did not observe spine shrinkage (Thomazeau et al., 2021; Vanderklish & Edelman, 2002) had low or undetectable levels of activity. None of those studies explicitly analyzed the levels of neuronal activity in their preparations, but those that detected spine shrinkage might in fact have been observing MIND-sLTD. For example, acute hippocampal slices used in Thomazeau et al. (2021) had their CA3 region cut out, leaving CA1 isolated, which might result in a lower degree of spontaneous endogenous activity, and thus, no appropriate conditions for MIND-sLTD. In contrast, the studies by Ramiro-Cortés and Israely (2013) and Sanderson et al. (2022) used slices with an intact CA3 region, which could drive higher endogenous activity in CA1. Moreover, Sanderson et al. (2022) were stimulating the slices with test pulses every 30 seconds and added picrotoxin to the ACSF, which could have induced enough circuit activity to promote MIND-sLTD (although they also used 2-chloroadenosine to reduce excitability). Strong support for this activity requirement comes from Ramiro-Cortés and Israely (2013), who blocked DHPG-induced shrinkage with TTX. The most recent study addressing this issue is Ramiro-Cortés et al. (2026), who have reported in a preprint an elegant protocol to induce single-spine mGluR-LTD using two-photon ppLFU, without using DHPG or TTX. With this method, they obtained evident long-term single-spine shrinkage (Ramiro-Cortés et al., 2026). The uncaging protocol itself could provide the glutamatergic signal needed to engage persistent remodeling, as we discuss below. Although they did not assess the effect of TTX, they performed the experiments using caged MNI-glutamate, which has previously been described to block GABAAR (Fino et al., 2009; Matsuzaki et al., 2010; Noguchi et al., 2011). Therefore, slices might have had elevated levels of neuronal activity in their experiments and thus MIND-sLTD might also be taking place. Interestingly, they blocked the persistent spine shrinkage with a protein synthesis inhibitor and partially with an NMDAR inhibitor, which is in agreement with what we have observed in MIND-sLTD (Fig. 3D-E).

### Molecular mechanisms underlying mGluR-induced neuronal activity-dependent spine shrinkage (MIND-sLTD)

The next question that arises is: what does neuronal activity provide that allows MIND-sLTD to occur? Increased circuit activity can engage several processes, including presynaptic glutamate release, postsynaptic depolarization, and signaling from neighboring cells such as astrocytes. Participation of other ligands or neurotransmitters simultaneously released, such as acetylcholine, epinephrine or serotonin, cannot be ruled out. Distinguishing among these possibilities will require additional experiments, but our single-spine experiments point to a predominantly local postsynaptic glutamatergic mechanism (Fig. 4). Although MIND-sLTD in bicuculline experiments was blocked by TTX (Fig. 3C), action potentials might not be necessary as the photoreleased glutamate in the synaptic cleft can produce the effect. Hence, a local glutamatergic signal may provide the activity-dependent component needed for MIND-sLTD to be initiated or expressed.

An interesting finding of our study is that mGluR activation and neuronal or synaptic activity must temporally coincide to trigger persistent spine shrinkage. Functional DHPG-LTD can be induced without concurrent synaptic stimulation (Huber et al., 2001), whereas our results show that structural DHPG-induced sLTD requires a temporal coincidence between mGluR activation and neuronal activity (Fig. 2). Single-spine experiments also suggest that there must probably be a spatial coincidence at the same synapses (Fig. 4). The next question, then, is what mechanism underlies this coincidence requirement. We do not currently know what synaptic mechanisms can explain MIND-sLTD, but we can think of several possibilities where mGluR and activity-induced signaling cascades converge on pathways controlling spine morphology through F-actin-regulatory factors, for example: activity-induced proteins such as Arc/Arg3.1, receptors such as NMDARs and AMPARs, actin-binding proteins such as cofilin, kinases such as p38-MAPK, and structural proteins such as synaptopodin. One of the possibilities that we addressed is the hypothesis that high levels of activity might activate immediate early genes producing plasticity-related proteins that could prime spine remodeling. Arc/Arg3.1 is particularly relevant due to its well-established role in functional mGluR-LTD in the hippocampus (Newpher et al., 2018; Park et al., 2008; Waung et al., 2008). We tested this hypothesis by extending bicuculline exposure from 1 h to 4 h to allow more time to synthesize and transport new activity-induced proteins (Fig. 2D). Longer pretreatment did not detectably increase shrinkage, and importantly, withdrawal of bicuculline before DHPG prevented spine shrinkage (Fig. 2E), suggesting that the accumulation of activity-induced proteins alone may be insufficient and that an additional signal must be available during mGluR activation. Thus, prior history of neuronal activity does not influence the extent of spine plasticity, but the temporal coincidence of activity and mGluR activation is critical to gate structural plasticity. Arc or other activity-regulated proteins could nevertheless participate if their translation, localization, or action is coupled to the concurrent activity. In fact, our experiments using cycloheximide (Fig. 3E) demonstrate that synthesis of new proteins is necessary for the maintenance of spine shrinkage, as also reported by (Ramiro-Cortés & Israely, 2013), although not necessary for the induction phase. It is especially interesting that MIND-sLTD shares this protein synthesis dependence with both functional mGluR-LTD (Huber et al., 2000) and structural NMDAR-sLTD (Thomazeau et al., 2021), both dependent on metabotropic signaling.

NMDARs were strong candidates to explain the coincidence effect, as they are known to play a crucial role in most forms of synaptic plasticity. They act as coincidence detectors of presynaptic and postsynaptic activity, bidirectionally governing the rules leading to both LTP and LTD (Malenka & Bear, 2004). They have also been strongly implicated in structural spine plasticity, also bidirectionally mediating the mechanisms of sLTP and sLTD (Kasai, Fukuda, et al., 2010; Stein et al., 2015). Depending on the Ca²⁺ influx through NMDARs or other Ca²⁺ channels, the NMDAR metabotropic signaling can induce either spine growth (NMDAR-sLTP) or spine shrinkage (NMDAR-sLTD) (Stein et al., 2021). However, persistent shrinkage remained detectable during AP5 treatment in our bicuculline experiments (Fig. 3D), and CPP did not block the spine shrinkage in our single-spine paradigm (Fig. 4E). These results suggest that ionotropic or metabotropic NMDAR activation is not the principal explanation for the activity requirement. NMDARs might still modulate the response: the smaller average shrinkage during AP5 treatment and the partial blockade with CPP reported by Ramiro-Cortés et al. (2026) are compatible with this possibility. MIND-sLTD may therefore engage alternative pathways or converge on some of the same downstream effectors while bypassing NMDAR activation.

These pathways could involve alternative sources of Ca²⁺ entry. We suggest a particularly plausible mechanism through which coincident activity could modulate the intracellular consequences of mGluR activation: a synergy between two Ca²⁺ signaling pathways. MIND-sLTD could be mediated by Ca²⁺ entry through voltage-gated Ca²⁺ channels (VGCCs) together with mGluR5-dependent Ca²⁺ signaling. Glutamate released during bicuculline-induced neuronal activity or locally by uncaging could activate AMPARs and kainate receptors (KARs), inducing depolarization that would open VGCCs. At the same time, activation of mGluR5 would trigger phospholipase C (PLC) signaling that leads to inositol-1,4,5-triphosphate (IP3)-mediated Ca²⁺ release from intracellular stores. In this model, AMPARs and KARs would play a critical role in the generation of MIND-sLTD. The downstream effector would be the special pattern of Ca²⁺ signal generated by both AMPAR-mediated depolarization and mGluR activation. In support of this model, Kim et al. (2015) showed that AMPAR-mediated depolarization facilitates mGluR5-dependent PLC signaling through Ca²⁺ entry via L-type VGCCs (Kim et al., 2015). It is known that both chemically induced LTD (via DHPG) and synaptic activity-induced LTD (via PP-LFS) trigger robust mGluR-dependent depression, but their intracellular mechanisms differ. PP-LFS-LTD depends on PLC and internal Ca²⁺ release (Bolshakov & Siegelbaum, 1994; Oliet et al., 1997), whereas DHPG-LTD does not (Mockett et al., 2011; Schnabel et al., 1999). Thus, just as distinct patterns of cytosolic Ca²⁺ can lead to opposite forms of NMDAR-dependent plasticity, namely LTP vs LTD (Malenka & Bear, 2004), or sLTP vs sLTD (Stein et al., 2021), it is possible that the combination of Ca²⁺ signals from mGluR-dependent cascades and activity-dependent cascades leads to two distinct forms of mGluR-LTD, one associated with spine shrinkage (MIND-sLTD) vs one without structural changes. This model predicts that PP-LFS-induced LTD may be more readily accompanied by spine shrinkage than DHPG-induced LTD under conditions of low activity. To our knowledge, spine structural plasticity has not been studied under electrophysiologically-induced PP-LFS-mGluR-LTD. The spine shrinkage observed after single-spine PP-LFU-induced mGluR-LTD in the recent preprint by Ramiro-Cortés et al. (2026) is consistent with this model. Our single-spine experiments also support this model, as we directly observed the effect of DHPG alone and DHPG concomitant with synaptic activity.

The pathways activated by mGluRs and neuronal activity may ultimately converge on the regulation of the actin cytoskeleton, which tightly controls spine morphology (Bosch & Hayashi, 2012). Rho-family GTPases, including Rac1, RhoA, and Cdc42, and actin-regulating proteins such as cofilin are plausible mediators of this convergence. Cofilin has been implicated in both spine enlargement (Bosch et al., 2014) and spine shrinkage (Hayama et al., 2013; Stein et al., 2020; Q. Zhou et al., 2004) after induction of NMDAR-LTP and NMDAR-LTD, respectively, and it is regulated by Ca²⁺ through the phosphatase Slingshot (Y. Wang et al., 2005). CaMKII is another attractive candidate to act as a central hub for MIND-sLTD as it links Ca²⁺ signaling to actin cytoskeleton reorganization and crucially participates in many forms of functional and structural plasticity (K. Okamoto et al., 2009), including spine growth (Matsuzaki et al., 2004) and spine shrinkage (Stein et al., 2020). There is evidence that both cofilin (Eales et al., 2014; Z. Zhou et al., 2011) and CaMKII (Mockett et al., 2011) participate in functional mGluR-LTD, but no study, to our knowledge, has addressed their role in structural mGluR-sLTD.

In our bicuculline-induced activity experiments, MIND-sLTD was blocked by TTX (Fig. 3C), which would suggest a role for presynaptic action potentials. However, glutamate uncaging experiments bypass the need for action potentials, as we provide the synaptic glutamate in the presence of TTX. Nevertheless, we cannot totally rule out the involvement of presynaptic (Nosyreva & Huber, 2005) or glial signaling. Postsynaptically generated endocannabinoids could retrogradely signal to presynaptic CB1 receptors and alter transmitter release, creating feedback between the shrinking spine and its presynaptic input. A relevant example is provided by Wu et al. (2024), who found that loss of synaptopodin altered functional mGluR-LTD, leaving a protein synthesis-independent component mediated by endocannabinoid signaling (Wu et al., 2024). The distinction between presynaptic and postsynaptic mGluR-LTD described by Sanderson et al. (2022) is another particularly relevant possibility. In their preparation, 30 µM DHPG recruited an NMDAR-dependent presynaptic form, whereas 100 µM DHPG could recruit an NMDAR-independent postsynaptic form, which was associated with spine shrinkage (Sanderson et al., 2022). If we assume that their slices had high levels of circuit activity (due to picrotoxin), their postsynaptic form of mGluR-LTD would correspond to our form of MIND-sLTD (we similarly used 100 µM DHPG in organotypic slices in bicuculline) while the presynaptic form would not produce enough mGluR activation to trigger the postsynaptic coincidence-sensitive system that translates into spine shrinkage.

### A role for the structural plasticity of spines

A conceptual implication of our study is that the structural plasticity of dendritic spines is governed by mechanisms that overlap with, but are distinct from, those governing functional plasticity, implying that structural plasticity plays a specific biological role other than accompanying functional changes. In the case of LTD, explicit or mechanistic dissociation between synaptic weakening and spine shrinkage has been demonstrated in NMDAR-LTD in rat hippocampus (X. Wang et al., 2007; Q. Zhou et al., 2004), in mGluR-LTD in mouse cerebellum (Sdrulla & Linden, 2007), and in mGluR-LTD in mouse and rat hippocampus (Thomazeau et al., 2021). Our findings identifying the temporal concurrence of mGluR activation and neuronal activity suggest that MIND-sLTD could provide a coincidence-detection mechanism through which local synaptic signals are integrated with the global activity of the surrounding network, determining which synaptic modifications acquire a lasting structural component and which do not. This structural change, therefore, represents an additional layer of regulation, where functional changes are consolidated depending on the integration of different signals. Spine elimination or synaptic uncoupling may constitute a further distinct layer of regulation (Stein & Zito, 2019), as loss of synaptic connection does not need to be preceded by spine shrinkage (Bastrikova et al., 2008). Thus, synaptic depression, spine shrinkage, and synaptic loss—like potentiation, spine enlargement, and spine formation—may represent connected but independent layers of metaplasticity that gradually consolidate the circuit remodeling that underlies processes like neuronal development, learning and memory (Hayashi & Majewska, 2005; Kasai, Fukuda, et al., 2010; Kasai, Hayama, et al., 2010). This capacity for selective structural remodeling may ultimately explain the biological advantage of organizing excitatory synapses on dendritic spines rather than directly on dendritic shafts.

## MATERIALS AND METHODS

### Animals

Sprague–Dawley (SD) rats of both sexes were used to prepare organotypic hippocampal slice cultures for experiments in Figs. 1–3. They were maintained and bred at the Animal Experimentation Unit of the Faculty of Pharmacy, Universitat de Barcelona, Spain. C57BL/6J mouse pups of both sexes were used to prepare organotypic hippocampal slice cultures for experiments in Fig. 4. They were maintained at the animal facility of the Center for Neuroscience, UC Davis, United States. All experimental procedures were performed in accordance with protocols approved by the corresponding institutional ethics committees and complied with European and United States regulations and guidelines for the care and use of laboratory animals.

### Organotypic hippocampal slice cultures and gene transfection

Hippocampal slices were obtained from rat or mouse pups at postnatal days 6-8 (P6-8), as described (Bosch et al., 2017; De Simoni & MY Yu, 2006; Stoppini et al., 1991). Slices of 400 μm thickness were plated in 6-well plates on a piece of semipermeable (0.4 μm pore size) BioporeTM interface membrane (Millipore, Cat. BGCM00010) positioned on top of a Millicell® Single-Well Standing Insert (Millipore, Cat. PICM0RG50). Slices were cultured at 35 °C in a humidified incubator with 5 % CO₂, fed in MEM (Minimum Essential Medium, Sigma-Aldrich, Cat. M4642) medium supplemented with 20% horse serum (Gibco, Cat. 11540636), 27 mM D-glucose, 6 mM NaHCO3, 2 mM CaCl2, 2 mM MgSO4, 30 mM HEPES, 0.01 % ascorbic acid and 1 µg/mL insulin (Sigma-Aldrich, Cat. I0516). The pH was adjusted to 7.3 and osmolality to 320-330 mOsm. The medium was changed every 2–3 days. Millicell inserts were reused after re-sterilization by sequential overnight washes in 100 % ethanol, 10% acetic acid, water, 100 % ethanol, followed by drying for 2 h under UV light.

Slices were biolistically transfected using a gene gun (Bio-Rad) at a helium pressure of 160 psi, after 5-7 days *in vitro* (DIV), as described (Woods & Zito, 2008). Slices used in Figs. 1-3 were transfected with plasmids expressing GCaMP6s under the CMV promoter (kind gift from Rachael Neve, MIT, Massachusetts General Hospital), and DsRed2 (Clontech) under the CAG promoter (from K. Futai, MIT, UMass Medical School), at a ratio of 7:3 (GCaMP6s:DsRed2). Slices used in Fig. 4 were transfected with a plasmid expressing EGFP under the CMV promoter.

### Time-lapse live fluorescence confocal microscopy

Time-lapse fluorescence imaging of living hippocampal slices was carried out using a Leica confocal microscope (DMI8) equipped with a 60 × 0.9 NA objective for spine morphology imaging or a 40× objective for calcium imaging. Transfected slices cultured for 11-15 DIV were transferred to the recording chamber and continuously perfused with carbogenated (95 % O2/5 % CO2) artificial cerebrospinal fluid (ACSF) at a flow rate of 3 mL/min and a temperature of 32-33 °C, maintained using an inline heater. ACSF contained 119 mM NaCl, 2.5 mM KCl, 26 mM NaHCO3, 1 mM NaH2PO4, 11 mM D-glucose, 2 mM CaCl2 and 0.2 mM MgCl2. GCaMP6s images were acquired at an excitation wavelength of 488 nm, while DsRed2 images were acquired at an excitation wavelength of 552 nm. The lowest light intensity sufficient to achieve an adequate signal- to-noise ratio was used to minimize photobleaching. For each slice, a well-isolated CA1 pyramidal neuron expressing both DsRed2 and GCaMP6s at moderate levels was selected, based on evident healthy dendritic morphology and no signs of fluorescent aggregates. In this neuron, time-lapse imaging was performed on a segment of a secondary or tertiary apical dendrite in the *stratum radiatum* as described (Bosch et al., 2014, 2017). Image z-stacks were acquired at 1-μm z-steps every 4 min for 92 min with a 3× digital zoom. Calcium imaging was performed before and after each experiment by sequentially imaging the entire field of view and then the soma of the selected neuron in a single optical section at 1-s intervals for 5 min. DHPG or control treatment was applied 30 min after the first image, i.e., after eight baseline images, by switching the perfusion input. After 10 min of treatment, perfusion input was switched back to DHPG-free ACSF until the end of the recording.

### mGluR-LTD induction and pharmacological reagents

mGluR-LTD was induced by applying 100 µM (R,S)-3,5-dihydroxyphenylglycine (DHPG, Tocris, #0342) for 10 min. DHPG powder was divided into aliquots and stored at −20 °C. Fresh 5 mM stock DHPG solutions were prepared in double-distilled H2O and used fresh or stored at −80 °C. In pharmacological experiments, slices were pre-incubated with the respective drug for 30 min before the beginning of the baseline period (i.e., 1 h before DHPG/vehicle application) and the drug was maintained in the bath throughout the entire experiment (except specific conditions in Fig. 2D-F). The following drugs were used at the indicated working concentrations: (-)-Bicuculline methiodide (Bic, 10 µM, Tocris, #2503), MTEP hydrochloride (1 µM, Tocris, #2921), LY367385 (50 µM, Tocris, #1237), D-(-)-2-Amino-5-phosphonopentanoic acid (AP5, 50 µM, Tocris, #0106), cycloheximide (CHX, 60 µM, Tocris, #0970), (R)-CPP (1 µM, Tocris, #0247), tetrodotoxin (TTX, 1 µM, Tocris, #1078), and 4-Methoxy-7-nitroindolinyl (MNI) caged L-glutamate (MNI-glutamate, 2.5 mM, HelloBio, #HB0423).

### Analysis of spine morphology and calcium signals from confocal time-lapse images

Spine morphology was analyzed using Fiji (ImageJ) software (Schindelin et al., 2012). Sum-intensity projections were generated from confocal z-stacks. Regions of interest (ROIs) of constant area were manually defined around 10–17 spine heads, four dendritic segments, and four background regions. Spines were selected if their entire morphology was clearly visible against a dark background, with clearly resolved heads and necks, throughout all frames of the time-lapse series, irrespective of their size and shape. Selected spines therefore encompassed a range of sizes and morphologies, including thin, thick, and mushroom-shaped spines, whereas dim filopodia and stubby spines were excluded. Spines were excluded if they overlapped with neighboring spines or other dendrites in any frame of the series. Small displacements of dendrites between time points were corrected using the StackReg plugin, and ROIs were manually repositioned when necessary. Spine volume was estimated from the mean gray value measured within each spine ROI. Background fluorescence was subtracted from spine measurements, and spine values were normalized to dendritic fluorescence to correct for global fluctuations in fluorescence intensity. Spine volume values were expressed as a percentage of the average baseline fluorescence intensity obtained before DHPG application.

Calcium imaging was analyzed using Fiji (ImageJ) software with ROIs comprising the entire image or the soma of the selected neuron. In each case, mean fluorescence intensity for each image in the time lapse series (consisting of 300 images: 5 min at 1 frame/second) was calculated and normalized to the intensity of the first image. Calcium peaks were identified using a custom-written algorithm as signals crossing an arbitrary threshold placed above noise.

### Two-photon imaging and single-spine glutamate uncaging

Organotypic mouse hippocampal slices cultured for 11-14 DIV were placed in the recording chamber at 28-29 °C and continuously perfused at 2 mL/min with carbogenated (95 % O2/5 % CO2) ACSF containing 127 mM NaCl, 2.5 mM KCl, 25 mM NaHCO3, 1.2 mM NaH2PO4, 25 mM D-glucose, 2 mM CaCl2, 0.2 mM MgCl2, 2.5 mM MNI-glutamate, and 1 µM TTX (∼310 mOsm, pH 7.2). MNI-glutamate was added at least 15 min prior to uncaging stimulation. EGFP-expressing CA1 pyramidal neurons (at 10-50 µm depth) were imaged using a two-photon microscope (Bruker) at 930 nm excitation wavelength with a custom perfusion system (Stein et al., 2019). For each neuron, image stacks (512 × 512 pixels, 1 µm z-steps) were acquired from a segment of secondary or tertiary basal dendrites at 5-min intervals.

Uncaging of MNI-glutamate at individual spines was performed after four baseline images using a parked beam (720 nm, 9–12 mW at the sample) at a point ∼0.5–1 μm from the spine head. Subthreshold low-frequency glutamate uncaging (subLFU) consisted of 60 pulses of 0.4 ms duration at 0.1 Hz. Spines between the 25th and 75th percentiles of the initial spine size distribution were selected to avoid unstable small spines and very large spines resistant to volume changes. In every experiment, we first confirmed the viability and capacity for structural plasticity of the selected CA1 neuron by inducing transient spine enlargement using a high-frequency uncaging protocol (HFU, consisting of 60 pulses of 2 ms duration at 2 Hz) in a distant segment of the same dendrite or another dendrite of the same neuron, located away from the subLFU spine site.

### Analysis of spine morphology from two-photon time-lapse images

Spine morphology was analyzed using custom-written MATLAB software. ROIs were manually defined around spine heads and adjacent dendritic segments in the optical section in which each spine appeared brightest and were adjusted, when necessary, at each time point. Integrated fluorescence intensities were measured from both spine and dendritic ROIs. Background fluorescence for dendritic ROIs was estimated from a distant ROI located away from labeled structures and subtracted from the integrated dendritic signal. For spine ROIs, background fluorescence was estimated from a neighboring ROI of identical dimensions positioned adjacent to the dendritic shaft and subtracted from the integrated spine signal to account for local background fluorescence. Dendritic fluorescence was monitored to verify its stability and was not used to normalize spine fluorescence. Spine volume was estimated from the resulting integrated fluorescence intensity values and expressed as a percentage of the average baseline fluorescence intensity obtained before stimulation.

### Statistical analysis

Data processing and statistical analysis were performed using Prism v10.0 (GraphPad Software). Data were normalized to baseline and are presented as the mean ± standard error of the mean (SEM). Short-term spine shrinkage (sSTD) and long-term spine shrinkage (sLTD) were calculated over the 6-18 min interval (4 time points) and the 46-62 min interval (last 5 time points) after the onset of DHPG/vehicle treatment, respectively, in Figs. 1–3, and the 10-15 min interval (2 time points) and the 30-40 min interval (last 3 time points) after subLFU stimulation onset, respectively, in Fig. 4. These values were statistically compared with values from the corresponding intervals in the control (mock or vehicle) condition. When compared to baseline, baseline was calculated as the interval from -18 to -2 min relative to DHPG/vehicle onset (5 time points) in Fig. 1–3, and the interval from -10 to 0 relative to subLFU stimulus onset (3 time points) in Fig.4. Outliers were identified using the ROUT method (Q =1 %) and removed from the data sets. Normal distribution was assessed using d’Agostino-Pearson or Shapiro-Wilk tests. If data were normally distributed, equality of variances was assessed using the F-test for two groups and the Brown-Forsythe test for three or more groups. For normally distributed data sets with equal variances, two independent groups were compared using a two-tailed unpaired t-test, paired measurements were compared using a two-tailed paired t-test, and three or more groups were compared using one-way ANOVA followed by Tukey’s, Dunnett’s, or Bonferroni’s multiple comparisons test. For normally distributed data sets with unequal variances, three or more groups were compared using Welch’s ANOVA followed by Dunnett’s T3 multiple comparisons test. For non-normally distributed data sets, three or more groups were compared using the Kruskal–Wallis test followed by Dunn’s multiple comparisons test. A statistically significant difference between groups was defined at the 95% confidence level (two-tailed), i.e., p < 0.05. Figure legends include information on the sample size, number of biological replicates, p value, and statistical test used for each experiment.

## ACKNOWLEDGMENTS

We are grateful to Maria Boerngen, Aina Asensio, Marta Pérez, Txell Martínez, Paula Rodríguez, Toni Luque, Núria Camarero, and Pau Gorostiza for advice, help and support. This research was funded by the Ministry of Science and Innovation grant PID2020-118817GB-I00 (SynWAVE) and in part by Fundació La Marató de TV3 (137/531-202224-30-31) to M.B., NIH R01NS137635 to K.Z., AEI/10.13039/501100011033 (PID2022-138105OB-C21 and CEX2021-001159_M Maria de Maeztu Unit of Excellence Program) to E.S., and PID2019-109285GA-I00, RYC2021-032313-I, PID2022-139822NB-I00, PCI2022-134995-2, Horizon ERC Consolidator grant 101088598 (HEU-101088598-DREAMNET) to A.R. E.R-F. is a recipient of an FPU fellowship (FPU22/00396) and a Joan Oró fellowship from AGAUR (2023-3 00065). N.S-R. is supported by an Investigo 2023 contract from the European Union NextGenerationEU and AGAUR.

## AUTHOR CONTRIBUTIONS

ERF performed most experiments and analyzed most data in Figs. 1–4, and contributed to writing the paper; NSR performed many experiments and analyzed data in Figs. 1–3; AS performed some experiments in Fig. 2; JF helped design and supervised Fig. 4 experiments; KZ helped design Fig. 4 experiments and contributed funding; KH and AP performed preliminary experiments; ES contributed resources and funding; AR contributed funding; MB conceived, designed and supervised the project, contributed funding, and wrote the paper.

## COMPETING INTERESTS

The authors declare no conflict of interest.

## Notes

### Competing Interest Statement

The authors have declared no competing interest.

